# Machine Learning Predictions Surpass Individual mRNAs as a Proxy of Single-cell Protein Expression

**DOI:** 10.1101/2024.12.11.627925

**Authors:** Josephine Fisher, Oliver Wood, Samuel Bullers, Lynne Murray, Li Li, Matthew A. Jackson-Wood

## Abstract

**Background:** Expansive repositories of single-cell RNA-seq data are now available. These data are often analysed assuming that mRNA abundance reflects the expression of their cognate proteins. However, post-transcriptional and translational regulation make mRNA an inadequate proxy for protein. High sparsity in low abundance mRNAs from single-cell transcriptomics data further complicates the extrapolation of protein expression levels. Although methods for single-cell surface protein quantification exist, they incur additional technical steps at greater expense and have yet to see wide-spread adoption. Computational approaches for protein imputation from scRNAseq data have been published, which learn transcriptome-wide patterns that predict protein expression. These models can then be applied to infer surface protein expression on RNA-seq only data, to increase the utility of existing data repositories.

**Results:** We tested 8 such methods and compared the accuracy of predictions between approaches, and against cognate mRNAs as a direct proxy. Predictions from the trained models outperformed the use of mRNA expression as a proxy. We identify notable cases of cell surface proteins with very poor correlation with their mRNA that were predicted very successfully by imputation using the whole transcriptome. We find cell type signatures are a major determinant of predicted protein levels and, as such, prediction methods require representative training data.

**Conclusions:** These results reiterate that mRNA level is not a reliable predictor of cell surface protein expression, and that whole-transcriptome informed imputation can improve protein estimations given appropriately trained models.

## Background

There has been widespread adoption of methods to profile transcriptomics at the single-cell level (scRNAseq) (1, 2). This has resulted in the publication of an abundance of scRNAseq data sets, including several high-profile projects to generate atlases of human health and disease (3–5). While scRNAseq is a powerful tool, it should be used with respect to its limitations. Limited read depth, transcript bias, and low capture efficiency can produce highly sparse scRNAseq data, with lower abundance genes often failing to be detected (6). Complexity of post-transcriptional and translational regulation means that mRNA transcript abundance may not reflect functional protein levels. The relationship between mRNA and protein expression could also differ by cell type and state.

New technologies seek to expand options for single cell profiling to measure multiple modalities simultaneously, some of which quantify protein expression alongside mRNA. These include CITE-seq stategies, such as AbSeq from BD Bioscience and Total-Seq from Biolegend, both antibody-based methods that can measure the expression of 100s of cell surface proteins (2, 7–10). In all three, oligonucleotide barcoded antibodies are used to bind and detect cell-surface protein. Subsequently the cell can be lysed, and protein abundance measured alongside the mRNAs by sequencing. Such data presents an opportunity to quantify the relationship between mRNA transcript quantities and surface protein expression. However, the relative novelty of these approaches and challenge of generating and profiling oligo conjugated antibody panels means that uptake remains comparatively limited. As an alternative to direct measurement a multimodal reference may be used to train a machine learning (ML) model and used to impute protein levels on the large volume of scRNA-seq data already available.

Various methods have recently been published to infer protein levels from scRNAseq using models trained on a multimodal reference. The package Seurat, commonly used for single-cell data processing, is capable of graph-based reference mapping to assign protein expression based upon cell-cell distances (11); sciPENN, totalVI, BABEL, scMMT and cTPnet all apply some variation of a neural network to generate predicted protein expression values (12–15); the method scLinear uses a comparatively simple approach, applying dimensional reduction followed by linear regression (16). A related unsupervised approach, SPECK, uses reduced rank reconstruction for protein prediction directly from RNA expression alone, clustering each gene separately without using a multimodal reference or whole-transcriptome context (17). To date there has been no independent comparison of the accuracy of these prediction methods and their value over using mRNA abundance alone as an estimator. Additionally, inter-method comparisons of performance when tested on different features and tissues are lacking. To address this, we present a comprehensive evaluation of 8 prediction methods, seeking to define their strengths and limitations.

Protein expression predictions were made on PBMC CITE-seq data from healthy donors, split into training and test partitions (11). We measured correlation between predicted and measured expression values and compared between methods and against the performance of the cognate mRNA as a proxy alone. We also evaluated the main factors driving predictive power by training and testing methods on cell type restricted partitions of the data and by testing the ability of models trained on healthy PBMCs to predict on unseen data from tumour tissue. Overall, we found transcriptome-wide predictions of cell surface protein expression outperformed using the proteins mRNA in isolation. Baseline expression level and variability between cell types were major determinants of a protein’s predictability. Our findings highlight that mRNA can be a poor representative of cell surface protein abundance, and that there is sufficient information in the wider transcriptome to make a more accurate inference of protein expression. However, current methods require a well-matched multi-modal reference, which may limit their applicability.

## Results

### Gene mRNA abundances are an inconsistent proxy of surface protein expression

Prior to testing prediction methods, we measured the naïve correlation between proteins and their mRNA abundances as a baseline expectation of correspondence. For this we utilised a large CITE-seq data set profiling PBMCs from healthy volunteers from a vaccine trial, published by Hao et al (11)(Fig. 1A). This has been used extensively in demonstrations of multimodal single cell analyses (18–22). It is comprised of 161764 cells, classified into 31 functional types. The diversity of cell types and protein features further supports generation of large independent training and test data sets in latter analyses.

**Figure 1.**
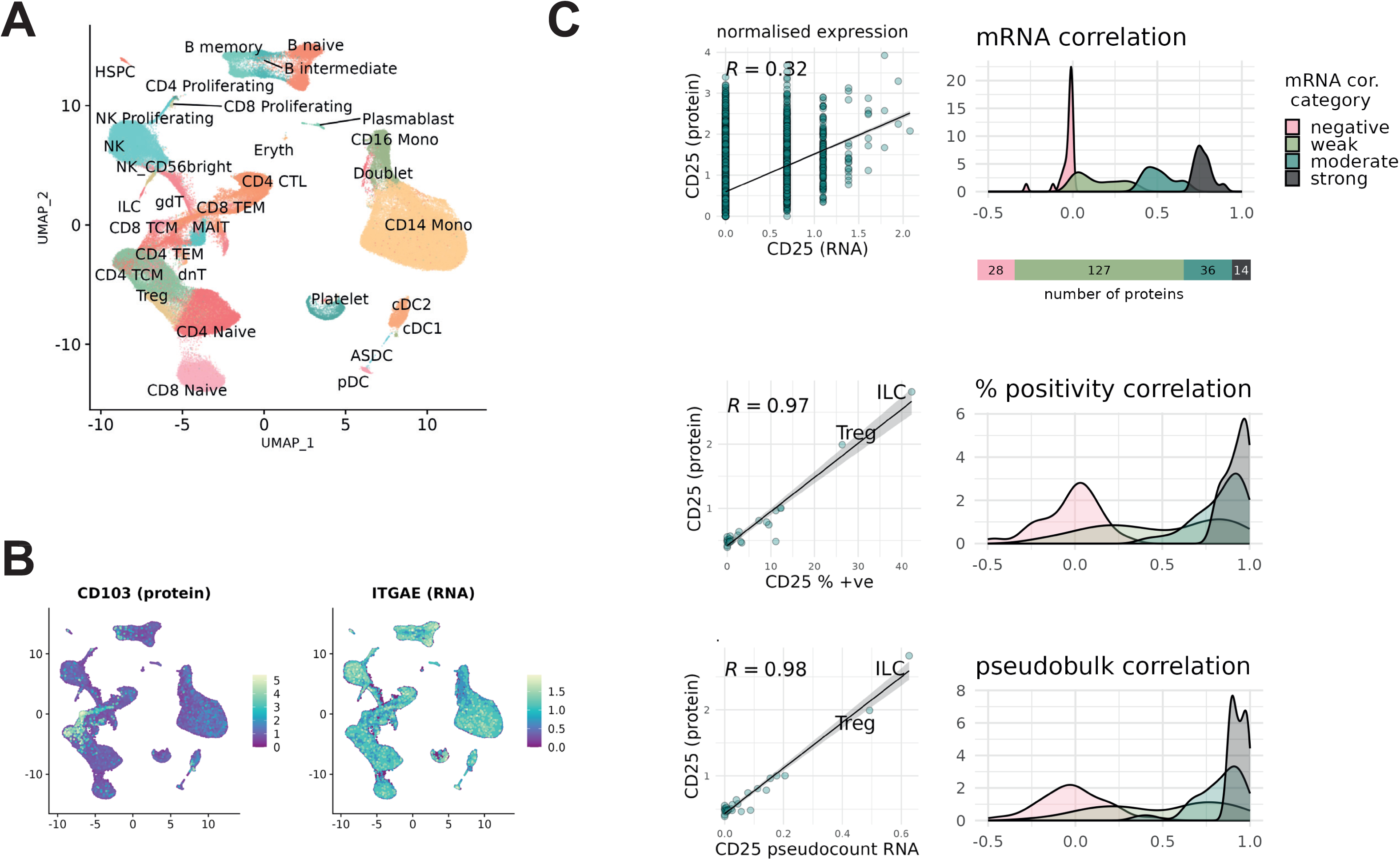
RNA-protein correlation in the Hao et al. CITE-seq data. A) UMAP showing the immune cell types profiled in the Hao et al. PBMC CITE-seq data. B) CD103 expression measured by normalised protein and normalised RNA. C) Density plots showing Pearson correlation of RNA-protein pairs measured by normalised RNA, proportion of RNA positive cells, and by RNA pseudobulk count. Features binned into correlation brackets according to correlation between normalised mRNA expression and measured protein.

Of the 228 proteins measured, 205 could be directly aligned with a single gene measured in the RNA data (excluding non-specific and control antibodies). We computed Pearson correlation between the measured protein and its corresponding mRNA (Supp. Fig. 1). We identified several cases where the expected expression profile as estimated by RNA was dramatically different from the expression observed by antibody measurement. A pronounced example was the tissue-residency marker CD103, showing extracellular protein detected only in a narrow subset of T-cells despite uniform abundance of RNA counts across all immune cell types (Fig. 1B). Of the 205 protein-mRNA pairs, 14 proteins have strong [0.7,1] Pearson correlation with normalised RNA abundance, 36 have moderate [0.4,0.7) correlation, 127 have weak [0,0.4) correlation, and 28 have negative correlation (Fig. 1C).

Proteins that are well approximated by RNA include several canonical immunophenotyping markers, such as CD3, CD16, CD14 and CD20. However, some markers of lymphocyte activation and cell state, such as CD25, showed poor correlation with RNA in circulating PBMCs (Fig. 1C). Some of these cases align with known biology; isoforms such as CD45RA and CD45RO are heavily controlled by post-transcriptional processes such as alternative splicing, so aren’t expected to align well with PTPRC expression(23). Other discrepancies have less obvious causes, and our results suggest a low probability that any given gene will have RNA measurements well-correlated with protein expression. To address this difficulty, we explore alternative metrics by which RNA can be used as an estimator for extracellular protein, computing either the percentage of RNA positive cells or the pseudobulk count over a cell-type (Fig1. C). Both approaches can recover a well-correlated, population-level estimate of median protein expression, generating similar results between the two methods. Examining changes in group distributions, we see these metrics can improve correlation values for strong and moderately correlated RNA-protein pairs. However, they largely fail to produce high correlation values for proteins with negative or low-valued correlation with the naïve mRNA proxy. Additionally, these metrics cannot be used where single-cell context is important and there remains a set of 102 proteins that fail to achieve correlation ≥ 0.7 using approximation by RNA counts, percentage positivity, or pseudobulk count (Supp. Fig. 2). CD103 is one such example, remaining resistant to these approaches due to fundamental disagreement between RNA and protein measurements.

### Cell surface protein prediction is comparable across most transcriptome wide modelling methods

Motivated by the insufficiency of protein estimation directly from source mRNA, we evaluated 8 published approaches for predicting protein abundances using the full transcriptomic profiles of cells. Again, performance was tested on the Hao et al CITE-seq dataset (11). We took a randomly sampled 20% fraction as the test (i.e. query) data and trained each method on the remaining 80%. Pearson correlation between the predicted and measured protein expression values was used as the primary evaluation metric, henceforth referred to as the prediction correlation.

All methods showed bimodality in prediction correlation across features, indicating a discrete set of features that are more challenging to impute than others (Fig. 2B). We sought to identify what characteristics might make a protein easier or more difficult to impute and found a linear association between the mean prediction correlation and total number of counts of a protein (Fig. 2C). Proteins with high standard deviation also trend towards having better average prediction correlation. Thus, features with strong and distinct signal in the data appear easier for prediction methods to impute.

**Figure 2.**
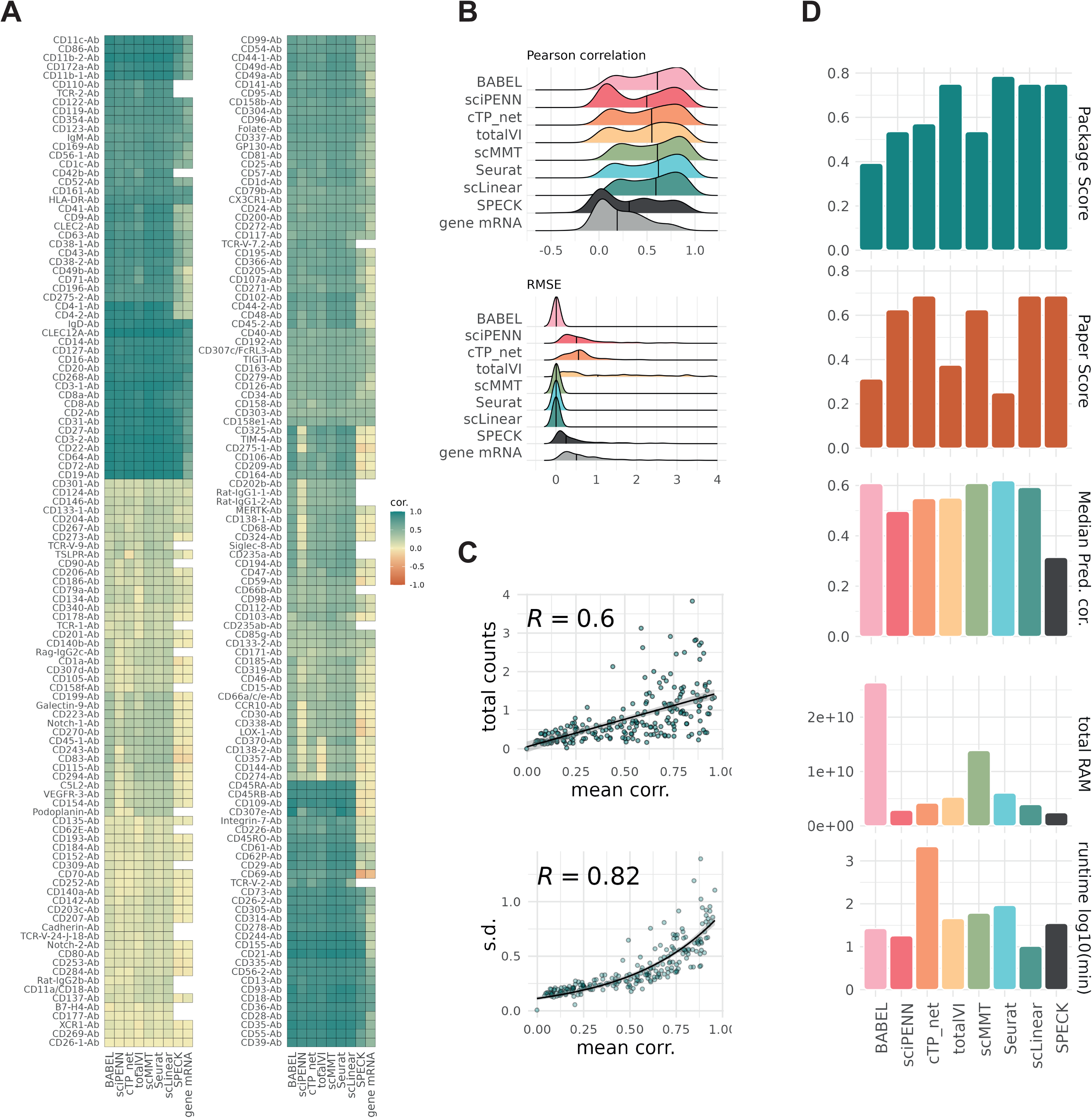
Prediction performance and method comparison. A) Heatmap showing Pearson correlation between predicted and measured protein expression values. B) Density plots showing distribution of Pearson correlation and RMSE for each method. C) Scatter plots showing average prediction correlation against total protein counts and standard deviation in the training data. D) Overview bar plots providing method summary, showing usability scores for the software and publication (1 being the highest), overall prediction correlation, RAM usage and runtime.

For many proteins there is low variability in prediction correlation across the different approaches, indicating a uniform ability across methods to impute expression (Fig 2A). A notable exception is the unsupervised strategy SPECK, which does not utilise protein data in its training. SPECK consistently produced predictions with lower correlation than other methods. In several cases SPECK produced near zero prediction correlation for features which were well-predicted by other methods, such as CD4 and CD9 (Fig 2A). SPECK also failed to produce a prediction for some features owing to a lack of RNA detection in the training data, likely due to dropout.

Most methods produced similarly correlated predictions and are not easy to distinguish by accuracy alone. We therefore additionally scored usability of the software and thoroughness of the associated publication, applying an adapted version of the framework presented in evaluations of single-cell integration and trajectory inference methods (24, 25). For comparison the scores are aggregated, with more detail given in Sup. Fig 3. All tested methods are described in peer reviewed journals, with software freely available from a version-controlled repository. Usage tutorials exist for all methods, although with varying levels of detail and relevance to protein imputation. With this context, an ideally performing method would produce high imputation accuracy, have a well-documented and accessible software package, and a thoroughly interrogative publication. These qualities vary across methods, and we were able to highlight discriminating points between them (Fig. 2D). BABEL is presented as being primarily designed for imputation of ATAC-seq peak data. The documentation is therefore opaque on how to implement protein prediction, and although the approach performs well it is challenging to use. In the cases of Seurat, totalVI and scMMT the publication is most concerned with validating alignment of modalities, with protein imputation left as an auxiliary application. The vignettes do detail the software usage well, so although the methods themselves aren’t deeply interrogated for efficacy they are simple to execute. Overall, in terms of usability and prediction outcomes, the best-scoring methods are cTPnet, sciPENN, scLinear and scMMT. These are accessible software packages that apply well-presented methods and achieve high prediction correlation values.

We also assessed the speed/scalability of all methods on ascending partitions of the data, running with access to identical computational resources. The data was split into 6.125%, 12.5%, 25%, 50% and 100% fractions, and methods were trained according to the previously described 80:20 training and test split. Median prediction correlation and RMSE over all features was measured, to identify any changes in performance corresponding to training data size. Resource usage and runtime were also evaluated. We found most methods produced predictions with correlation stable to changes in training data size (Supp. Fig. 4). An exception is scMMT, which showed dramatically reduced performance at data size lower than 50% (approx. 80K cells). Regarding resource usage, we find neural-net based methods to be most demanding. BABEL and scMMT used the most RAM during training (Fig. 2D). The five neural-net based methods were also the highest consumers of CPU resources. cTPnet runtime appears to scale particularly poorly with size of training data, although this may be possible to mitigate by adjusting the training patience parameter.

### Whole transcriptome gene expression predicts protein abundance better than source mRNA alone

Across all tested methods there were many proteins that consistently showed improved prediction correlation with genome wide imputation compared to direct estimation from cognate mRNA (Fig. 3A). Overall, ML imputation maintains comparable median prediction correlation to aggregate RNA metrics and produces no negative correlation values. Of the 205 proteins for which cognate RNA data is available, 70 fail to achieve moderate prediction correlation >0.4 by either direct RNA estimation or aggregate RNA measures (Supp. Table 1); 23 of these achieve mean prediction correlation >0.4 when using ML methods.

**Figure 3.**
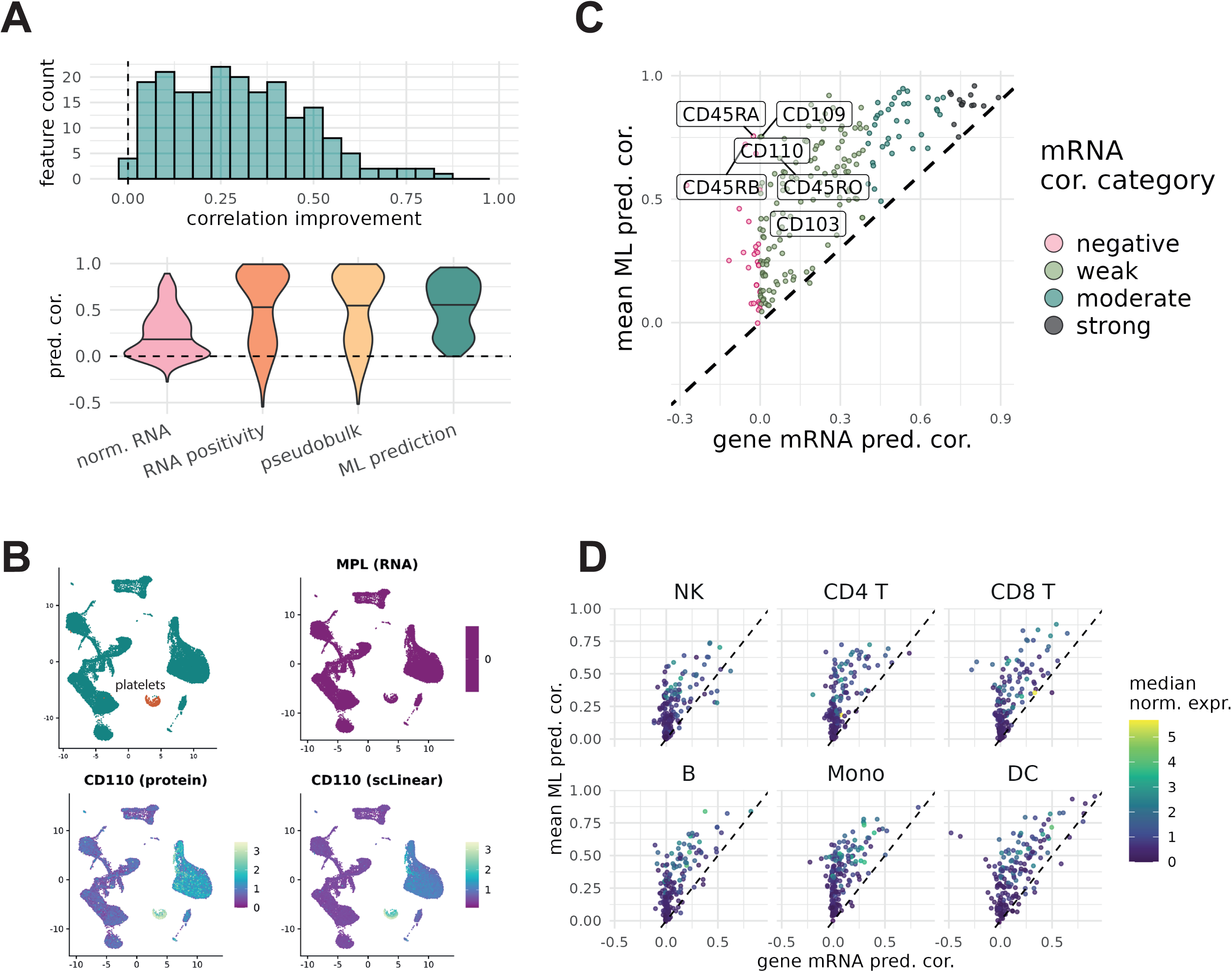
Improvement in prediction correlation using ML methods. A) Histogram of difference in correlation coefficient between mean prediction correlation and gene mRNA correlation. B) UMAP panels highlighting platelet population and showing MPL (CD110) RNA expression against protein expression and scLinear predicted values. C) Gene mRNA correlation plotted against mean prediction correlation across methods, coloured by correlation categories defined in Fig. 1C. Features with large correlation improvement are annotated. D) Correlation scatter as in C), split by cell type.

The utility of protein imputation is highlighted by the feature CD110, a thrombopoietin receptor commonly detected on platelets (1). Platelets contain no nucleus, and typically have only a small amount of legacy mRNA transferred from megakaryocytes (26). The CD110 transcript MPL is undetected in the Hao et al. test data rendering direct estimation impossible, however CD110 protein is enriched on the platelet surface. Using the full, albeit limited, landscape of platelet RNA, ML prediction methods can learn the transcriptional signature of platelets and recovers CD110 enrichment in the population (Fig. 3B). As a result, average prediction correlation across methods for CD110 is strong, at 0.75. Other examples benefiting from imputation include protein isoforms, which show a marked improvement in correlation using ML methods. Both CD45RA and CD45RO are predicted with much greater accuracy by the tested methods than by estimation from PTPRC mRNA alone, achieving mean correlation values of 0.76 and 0.67 respectively (Fig. 3C). Proteins that are well correlated with mRNA abundance such as CD20 and CD16 remain accurately predicted by ML methods, with some also showing a slight improvement in correlation. Splitting the data by cell type there were a small number of features that were better represented by mRNA than predictions (the most in dendritic cells) but for the majority of features the predictions improved upon mRNA in all cell types (Fig. 3D).

Only one protein showed reduced average prediction correlation with ML methods (CD301 reduced by 0.025). A subset of 70 proteins were poorly estimated by both normalised RNA-proxy and ML methods having prediction correlation <0.4 in both cases (Supp. Tab. 1). In some cases, these low values are expected; secreted proteins such as galectin-9 are unlikely to be well-captured by CITE-seq measurements targeting the cell-surface. Other weaker predictions may be due to low abundance or low variability of the proteins in the training data (Supp. Tab. 3). Overall, our results show ML imputation to either match or improve upon the naïve prediction correlation produced by normalised mRNA expression.

### Prediction accuracy depends strongly upon transcriptional signatures of cell type

Given that the proteins best predicted were often the most variable across the data set we investigated if cell type, a likely prominent source of variation, was an influential factor in ML inference. As an initial indicator of this we identified specific examples where the high correlation of prediction was absent when looking within an individual cell type compared to across all cells (Fig. 4A); i.e. methods may be good at predicting high level differences in protein expression between cell types but poor at quantifying the finer resolution differences in expression between very similar cells. Indeed, when we compared the prediction correlation as measured across all cells to that within individual cell types across all proteins, we noted a pronounced and consistent drop in accuracy (Fig. 4B).

**Figure 4.**
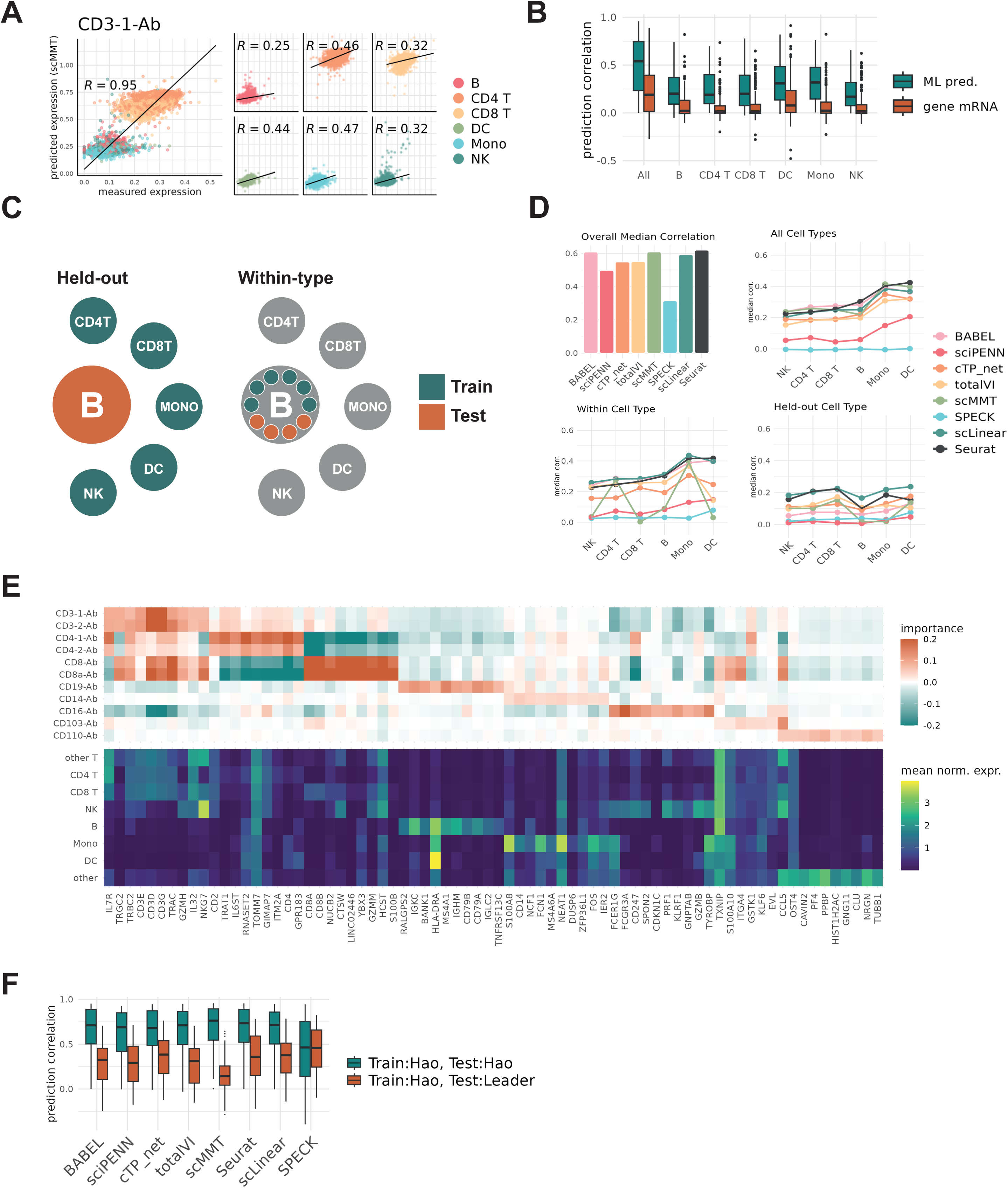
Sensitivity of prediction performance to training data composition. A) Scatter plot showing correlation of scMMT prediction with measurements, computed over all cell and over cell-subtypes. B) Boxplot showing mean prediction correlation over methods, compared against gene mRNA correlation. Correlation values measured over all cells, and for each cell type individually. C) Schematic of training and test sets in investigation of cell type sensitivity. D) Panels showing bar plot of overall prediction correlation across all cells, alongside median prediction correlation in each cell-type individually (All Cell Types), when only one cell type is used for both testing and training (Within Cell Type), and when a cell type is held out as the test set (Held-out Cell Type). E) Heatmap of top 10 most important RNA features in the scLinear prediction model, shown for a selection of proteins. F) Boxplot comparing prediction correlation in Hao et al. data against prediction correlation with methods trained on Hao et al and tested on Leader et al.

To interrogate this further, we ran protein imputation on two variations of the Hao et al. data (Fig. 4C). In the first, each cell type was isolated and used for both training and test partitions as before. This was intended to explore changes in performance with reduced phenotypic variability in the training data. In the second variation, one cell type was withheld as the test set and the prediction models were trained on the remaining cells. This tested the methods ability to infer protein expression on unseen cell types. We refer to the two training variations as within-cell type and held-out cell type respectively. We compared them to the all-cell case, where the entire data was used in both testing and training (Fig 4D).

In the within-cell type variant we saw mostly similar prediction correlations as in the all-cells case, with only minor reductions in median prediction correlation for most methods. However, scMMT showed pronounced sensitivity to the restriction of cell type information, producing substantially lower prediction correlation in NK cells, CD8 T cells, B cells and dendritic cells. In the held-out variant, all methods exhibit uniform decline in prediction correlation. When imputing protein on unseen cell types, no methods achieve median prediction correlation exceeding 0.25 across all features. Taken together, our results demonstrate a pronounced sensitivity of reference-based prediction to the composition of the training data.

To further investigate the cell type dependence of predictions we explored the features determining model predictions. Given the black-box nature of most of the methods, we focused on scLinear which provides measures of feature importance for its model. We plotted the top 10 most important features predicting a selection of cell type marker proteins and noted distinct mRNAs were important for the prediction of each (Fig. 4E). Some of these were correlated cell type markers that might be expected. For instance, expression of the B-cell marker CD19 was primarily determined by expression of other canonical B-cells markers such as MS4A1 and CD79. The T-cell markers CD3, CD4 and CD8 were strongly informed by their corresponding mRNAs and other genes predominantly expressed in T-cells. However, we also observed a small selection of non-obvious genes with high importance for protein imputation but also broad, cross-phenotype, expression. This suggests potential interactions with other genes in the model adding to predictive information. We also observed feature importances aligning with expected biology. For example, protein expression of the tissue residency marker CD103 is informed by a collection of mRNA features with related function: the genes EVL and ITGA4 both have established relevance to immune tissue infiltration (27, 28), and CCL5 is understood to recruit peripheral immune cells to inflammatory sites (29, 30).

As a final test of the ability to predict on unseen cell/tissue types, we tested protein prediction using the models trained on PBMC data on scRNAseq data from a non-small cell lung cancer CITEseq data set(31). We found that overall prediction correlation was substantially lower for all reference-based methods, but SPECK maintained a stable but lower median value (Fig. 4F). These results may indicate overfitting of the models to cell type characteristics in the PBMC training data, producing poor accuracy when tested on data of a different biological context.

## Discussion

Our exploration of CITE-seq data confirms that RNA does not always provide an accurate proxy measure for extracellular protein. Many post-transcriptional and post-translational mechanisms could explain the mismatch between intracellular RNA, and protein at the cell surface. Alternative splicing, intracellular trafficking apparatus, protein-protein interaction and decay/secretion rates all influence the quantity of protein maintained on the cell surface (32). This complex chain of events between RNA and protein measurements makes it difficult to determine how any single protein relates to its corresponding RNA. One can instead use an aggregate measure over a population of interest. We found that RNA positivity and pseudobulk expression both produce very similar outcomes, and often provide estimates well-correlated with median protein expression in the target population. However, these summary metrics remove the single-cell granularity of data and will still perform poorly where there is a biological disconnect between mRNA and protein, as with protein isoforms. Our comparison of ML methods suggests that whole transcriptome-based prediction provides a viable alternative to protein estimation and can recover obscured RNA-protein relationships through an understanding of cellular phenotype. Caveats remain regarding the selection of suitable training data and poor performance on a fraction of protein features.

The 8 tested ML prediction methods all largely produced similar results, leaving little room for discrimination by accuracy. The exception is SPECK, the only method not to utilise a protein reference. SPECK produced relatively poor prediction correlations in all tests, further emphasizing the limitations of inference directly from sparse RNA data where the target transcript may be completely undetected. The remaining 7 approaches are supervised methods and use a multi-modal protein/mRNA training reference. All produced predictions well-correlated with measurements, with values consistently better than mRNA alone. Regrettably all 7 methods also appear to suffer from overfitting to the PBMC reference, showing reduced performance when tested on tissue data unseen in training. The main differentiating factors between methods were runtime, resource usage, and sensitivity to scale and composition of training data. scLinear and Seurat both maintained performance stable to change in size of the training data, and both methods complete quickly without high computational resource requirements. Of the available options, we recommend these methods as accessible and capable strategies for protein imputation.

In our analyses prediction correlations varied strongly across proteins, with individual features having uniformly strong or weak correlations over methods. This suggested some inherent qualities of certain proteins could cause to them be challenging to accurately impute. We found that ML-prediction is not a one-size-fits-all solution for any target. Prediction methods tend to maintain good performance on cell-type defining features already well-approximated by RNA expression, such as CD3 and CD4. They can produce modest improvement in prediction correlation in many other cases with poor mRNA correlation and can rescue intractable features for which RNA is entirely undetected. However, some key immunoregulatory proteins remain poorly represented by both cognate mRNA and prediction methods. The checkpoint receptors BTLA (CD272), TIGIT and CTLA4 all failed to achieve strong average prediction correlation (>0.7) using ML methods but did pass that threshold by RNA aggregation metrics (positivity, pseudobulk) (Supp. Table 1). In such cases we find no reliable, single-cell resolution, estimation strategy. One is forced either to use aggregate metrics or measure the protein expression directly using a method such as CITE-seq.

We saw reduced performance when training and testing on different cell type subsets of the PBMC data, and when applying PBMC trained models to predict expression on unseen lung/tumour tissue. This implicated dataset-specific cell type transcriptional signatures as strong determinants of model performance, rather than nuanced learning of consistently relevant translational regulators. Feature importance in scLinear prediction supported this, revealing the genes with most predictive power to have highly differentiated patterns of expression by cell-type. We also observed well-predicted proteins to be those with high counts and variability in the data, generally having a strongly discriminated pattern of expression across cell types. These outcomes suggest a need for training and query data to be of similar phenotypic composition and with similar characteristics of gene expression, to avoid mismatch and misinterpretation of phenotype signatures. This may limit applicability of these methods where well-defined CITE-seq data are not available. Even in such instances it may be possible to design efficient, cost-effective hybrid studies. One could apply CITE-seq to a small subset of samples, and then use ML-imputation to infer protein on higher-throughput, highly multiplexed RNA data that can be generated more affordably (33–35).

Accurate quantification by CITE-seq can be challenging in the absence of high quality, validated, antibodies. In such instances, alternate technologies such as mass spectrometry could provide an alternative solution to measure protein without reliance on antibodies. Recent advancements in sample preparation and equipment sensitivity have enabled sensitive single-cell mass spectrometry albeit at a much lower throughput in terms of cell count (36). Going further, a range of new single-cell technologies expanding upon CITE-seq aim to profile intracellular protein and post-transcriptional features, offering new ways to map the pathways between transcription and extracellular protein(37–39). This increasing range of experimental and bioinformatic tools to study protein expression will further reduce the reliance on transcriptomic data as a proxy and could similarly be considered for training models for transfer learning.

### Conclusions

Our study highlights the inadequacy of mRNA abundance as a proxy of protein expression in single-cell data. We have evaluated the utility of transcriptome-wide ML based protein prediction as an alternative approach and define the limitations of these methods. Accurate estimation of cell surface protein expression is of great importance for the definition of cell types, study of disease pathology, and identification of therapeutic targets. Thus, adoption of models to more accurately estimate cell surface protein expression could prove valuable to improve the utility of large existing repositories of scRNA-seq data. Focus should be given to identifying and generating suitable reference data sets to learn from and the development of more advanced, globally applicable, foundational models associating cell transcriptomes with cell surface protein measures.

## Methods

### Data and preprocessing

We explored performance of protein imputation methods on two CITE-seq datasets. The first is the Hao et al. PBMC data (GSE164378[https://www.ncbi.nlm.nih.gov/geo/query/acc.cgi?acc=GSE164378]). This dataset describes blood samples obtained from eight volunteers enrolled in an HIV vaccine trial, taken at baseline and at days 3 and 7 following vaccine administration. Whole transcriptome RNA profiles are measured on 161,764 cells. A panel of 228 antibodies was used to profile surface proteins on all cells. This data was downloaded as a Seurat object provided by the New York Genome Center [https://atlas.fredhutch.org/nygc/multimodal-pbmc/], used with no further processing or cell filtering. We retained the log-normalized RNA counts and CLR normalized protein counts. The second CITE-seq dataset used is the non-small cell lung cancer data presented by Leader et al. RNA counts are provided under accession GSE154826[https://www.ncbi.nlm.nih.gov/geo/query/acc.cgi?acc=GSE154826], and protein data is available via the github repository [https://github.com/effiken/Leader_et_al]. The data contains RNA profiles for 361,929 cells, taken from tumour and non-involved human lung tissue. Of these, protein was measured on a subset of 230,820 cells. For our analyses we used protein data for 82 antibodies, taken from distinct panels measured on different patient subsets. For any protein-specific analyses (correlation computation, etc.) we include only cells on which the protein was measured. To align with our other analyses, we applied CLR normalization to the protein counts in this dataset.

### Aggregate RNA metrics

In our evaluation of correspondence between RNA and protein, we explore the value of using percent positivity and pseudocount estimators. Percent positivity is computed simply as the proportion of cells in a group in which the transcript was detected. Pseudobulk counts were computed using the Seurat function *AggregateExpession().* RNA counts were summed over donor and cell-type. The final pseudobulk value per cell-type was taken as the mean value over donors.

### Protein Imputation

We ran 8 protein prediction methods using a custom Nextflow pipeline [link to code repository when available] (40). Code for model training and prediction was sourced from author-provided github repositories or vignettes for each method. Arguments controlling prediction strategy were set to default according to vignettes or example code. Prior to prediction the data was partitioned into a training and test set, taking an 80:20 split, selecting cells at random. Predictions methods were trained and tested on the same partitions. Predictions made on the test data were evaluated against the measured protein by computation of Pearson correlation and RMSE. All imputation methods were left to apply built-in normalization as defined in the published software, and were provided unnormalized RNA counts as input.

### Evaluating Prediction Correlation

When evaluating baseline correlation between mRNA and protein in the Hao et al. data, we first had to determine suitable alignment between RNA and protein features. Where antibodies had no sensible, direct mRNA equivalent (isotype controls, TCR binding antibodies) they were dropped from correlation analysis. Where correlation was not possible to evaluate due to high sparsity and RNA dropout, we set prediction value to zero to facilitate comparison with other methods. RNA-protein correlations were computed only on cells from the test set, for parity with ML-prediction correlations. Where prediction correlation is evaluated over a set of features, we take the median correlation value. Where prediction correlation is evaluated over ML methods we take the mean correlation value.

### Prediction Benchmarking

To assess comparative performance of prediction methods Nextflow configuration was used to ensure each prediction process had access to identical memory and CPU resources, fixed at 240GB and 32 cores. The trace and execution reports generated by Nextflow were used to report runtime and resource usage for each method.

To explore how prediction results change with differences in training data, we ran the methods under three separate variations.

1. The data was subsampled into 6.125%, 12.5%, 25%, 50% and 100% fractions. Following this, it was split into a training and test set in an 80:20 ratio. Prediction methods were then trained and tested. This approach was intended to measure differences in performance as the size of the training sample increases.
2. For each cell type group in the set B, CD4 T, CD8 T, NK, Mono and DC the selected type was entirely held out as the test set. Methods were trained on the remaining cells, and prediction correlation measured on the held-out cells. This approach is intended to assess ability to accurate predict protein on unseen cell types.
3. For each cell type group in the set B, CD4 T, CD8 T, NK, Mono and DC the selected type was taken as the entire data and split into a training and test set in an 80:20 ratio. All other cells were discounted. Prediction methods were trained and tested on only one cell type at a time. This method was intended to measure capacity to detect within-cell type variation in protein expression.

### Feature Importance in scLinear

While not possible for all prediction methods, we were able to extract a measure of feature importance from the linear models generated by scLinear. As described by the package authors, feature importance is calculated as the Jacobian matrix of the predicted ADT values, with respect to the input RNA matrix. This is computed as the product of the Jacobians for the singular value decomposition, linear regression and z-score normalisation components of the model. Further details of computation are given by Hanhart et al. (16). Genes with large, positive importance value are understood to be influential over prediction outcomes.

### Usability Scoring

We measured usability with an adapted version of a framework previously applied to benchmark methods for single-cell integration and pseudotime trajectory inference (24, 25). Methods were evaluated according to 11 criteria (Supp. Table 2) which measure both how accessible the software is to use, and how thoroughly the publication evaluates prediction accuracy against other approaches. The first two criteria are simply that the method be available on a version-controlled repository, and that the underlying code is freely available and executable on an open-source platform. Four conditions are used to evaluate the quality of available software tutorials. These are that a tutorial be freely available, that it shows the imputation method in different scenarios, that it shows how to run the method in a non-native language, and that running the tutorial produces no notable errors. The documentation of function inputs, outputs and intended use is also scored. Finally, a set of four criteria evaluate the publication. The first is simply that the method is published in a peer reviewed journal. The others assess the number of datasets on which the method is tested, the diversity of robustness assessments, and breadth of comparison with other protein imputation strategies.

## Declarations

### Availability of data and materials

Two published CITE-seq datasets were used to evaluate imputation method performance. The first is the Hao et al. PBMC data (GSE164378[https://www.ncbi.nlm.nih.gov/geo/query/acc.cgi?acc=GSE164378]). The second is non-small cell lung cancer data presented by Leader et al. RNA counts are provided under accession GSE154826[https://www.ncbi.nlm.nih.gov/geo/query/acc.cgi?acc=GSE154826], and protein data is available via the github repository [https://github.com/effiken/Leader_et_al]. The Nextflow pipeline and accompanying scripts used for method execution and evaluation are available in the associated github repository [https://github.com/fisherj2/protein_imputation].

### Contributions

J.F. and M.JW were responsible for design of analyses and manuscript preparation. J.F. conducted computational work and figure compilation. S.B., O.W., L.M. and L.L. contributed to data interpretation and manuscript revision.

## Acknowledgements

Christopher Paluch

---

**Supplementary Figure 1. The correlation between protein and RNA measurements.** Bar plot showing Pearson correlation between normalised mRNA and measured extracellular protein, for 205 features. Bar coloured according to value.

**Supplementary Figure 2. Comparison of prediction comparison by different approaches.** Heatmap of prediction correlation values by estimation approach, with mean value taken over ML methods.

**Supplementary Figure 3. Evaluation of predictions methods by usability scoring.** Method usability metrics broken down by category, with 1 as the highest score and 0 as the lowest.

**Supplementary Figure 4. Performance and resource usage of prediction methods trained on ascending fractions of data.** A) Line plots showing median prediction correlation and RMSE for each method, as the data size is varied over subsamples. B) Boxplots showing resource usage and runtime for each method, over subsamples of the data.

