## Supplementary Figure 1 for "Machine Learning Predictions Surpass Individual mRNAs as a Proxy of Single-cell Protein Expression"

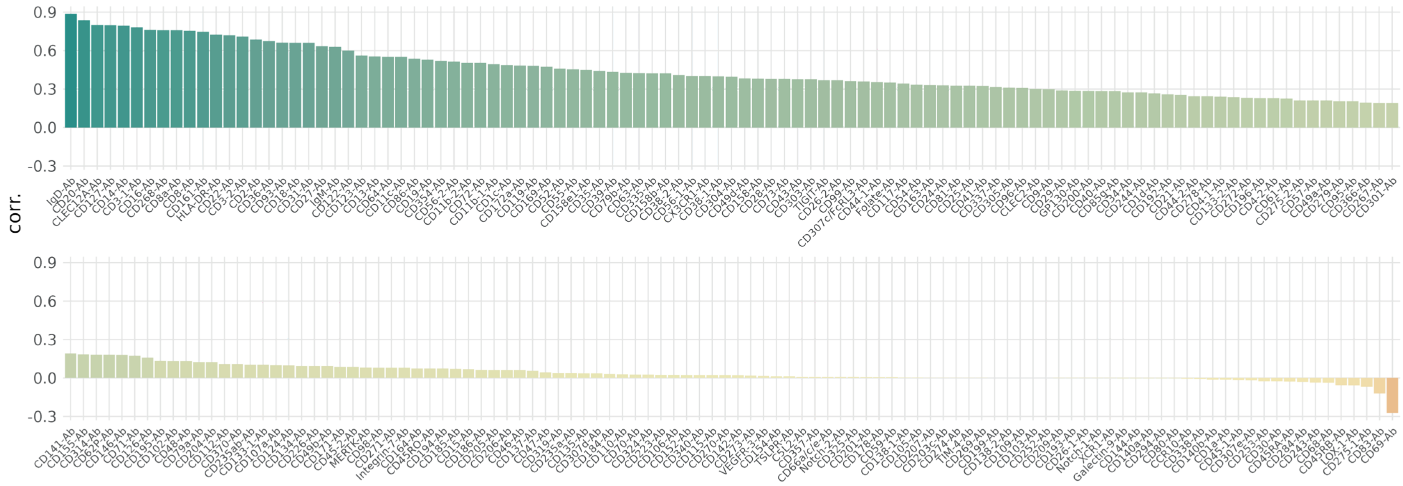


**Supplementary Figure 1. The correlation between protein and RNA measurements.** Bar plot showing Pearson correlation between normalised mRNA and measured extracellular protein, for 205 features. Bar coloured according to value.
