## Supplementary Figure 2 for "Machine Learning Predictions Surpass Individual mRNAs as a Proxy of Single-cell Protein Expression"

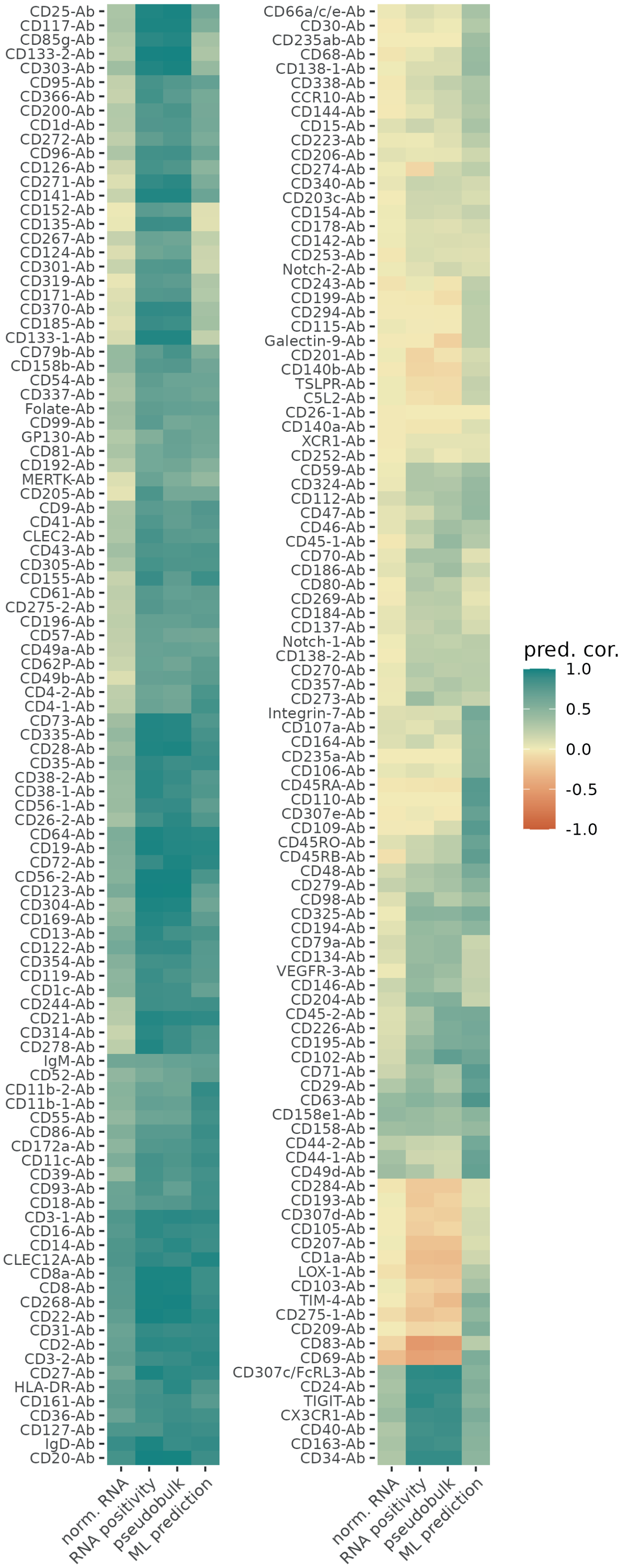


**Supplementary Figure 2. Comparison of prediction comparison by different approaches.** Heatmap of prediction correlation values by estimation approach, with mean value taken over ML methods.
