## Supplementary Figure 3 for "Machine Learning Predictions Surpass Individual mRNAs as a Proxy of Single-cell Protein Expression"

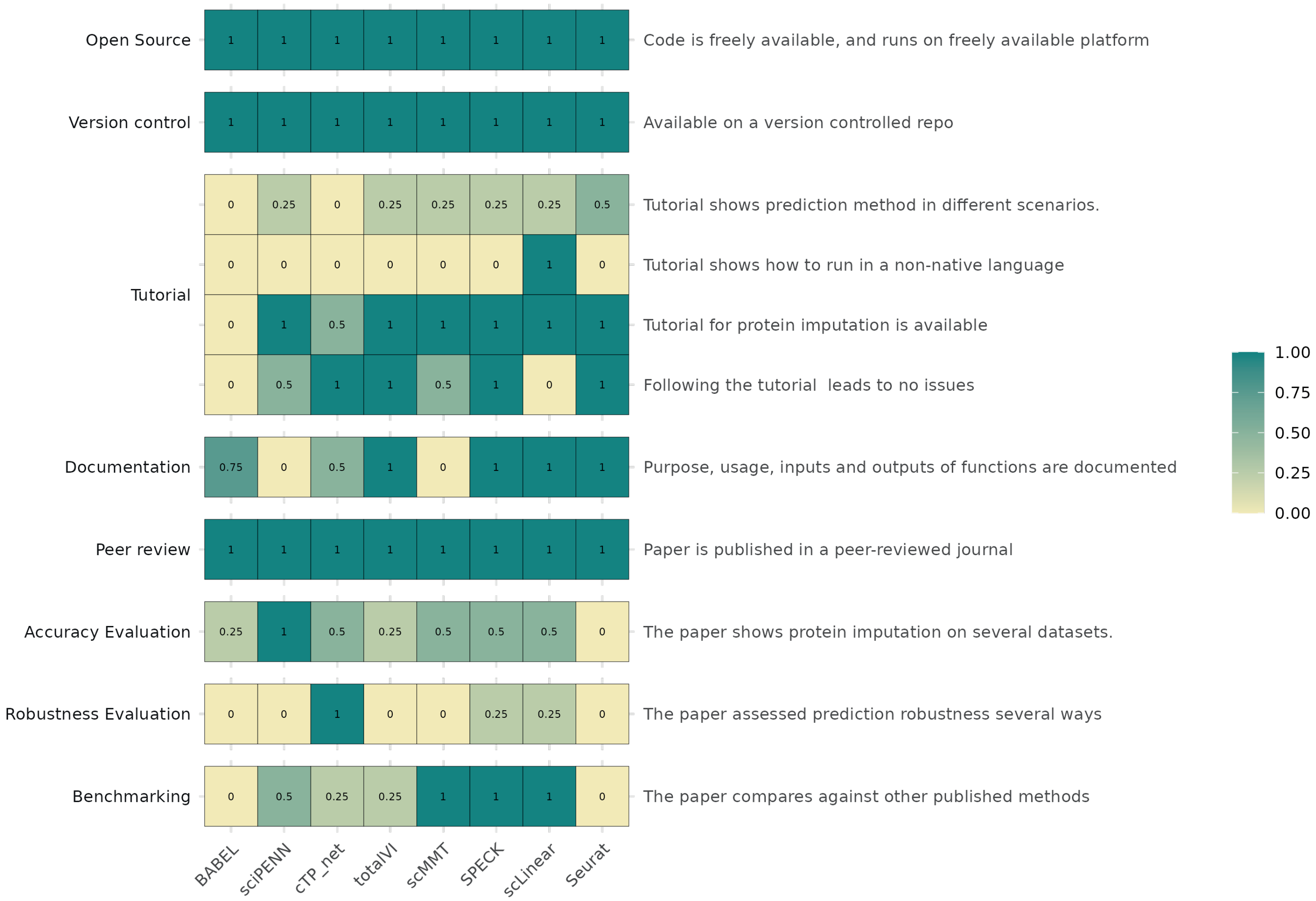


**Supplementary Figure 3. Evaluation of predictions methods by usability scoring.** Method usability metrics broken down by category, with 1 as the highest score and 0 as the lowest.
