## Supplementary Figure 4 for "Machine Learning Predictions Surpass Individual mRNAs as a Proxy of Single-cell Protein Expression"

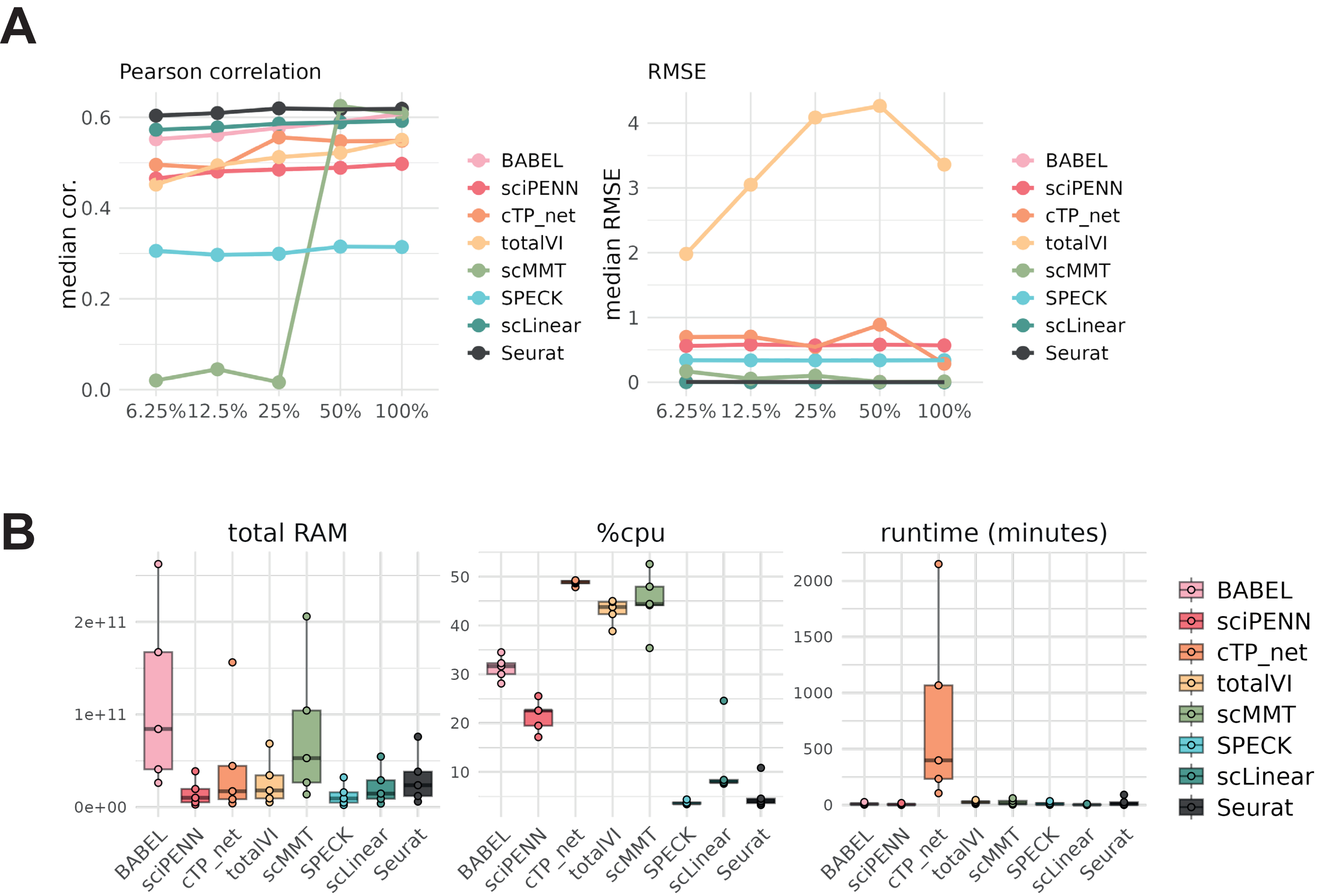


**Supplementary Figure 4. Performance and resource usage of prediction methods trained on ascending fractions of data.** A) Line plots showing median prediction correlation and RMSE for each method, as the data size is varied over subsamples. B) Boxplots showing resource usage and runtime for each method, over subsamples of the data.
